# A Minimal Model Explains Aging Regimes and Guides Intervention Strategies

**DOI:** 10.1101/2025.08.25.671954

**Authors:** Peter Fedichev, Jan Gruber

**Affiliations:** GERO PTE. LTD., 133 Cecil Street 14-01 Keck Seng Tower, Singapore 069535; 1 Healthy Longevity Translational Research Program, Yong Loo Lin School of Medicine, National University of Singapore, Singapore; Department of Biochemistry, Yong Loo Lin School of Medicine, National University of Singapore, Singapore

## Abstract

Aging varies widely across species yet converges on universal laws such as Gompertzian mortality. We propose a minimal framework that reduces this complexity to three macroscopic variables: the leading stress response (*z*_0_), cumulative entropic damage (*Z*), and regulatory noise strength (*D*_0_). The model predicts two regimes. In unstable species such as flies and mice, intrinsic instability drives exponential divergence of biomarkers and mortality. In stable species such as humans, linear damage accumulation steadily erodes the leading stress response, producing a hyperbolic trajectory toward a finite maximum lifespan. The framework reproduces survival curves and methylation dynamics across taxa and organizes intervention strategies into three levels: modulating stress responses, reducing noise, and slowing entropic damage, offering a roadmap for extending human healthspan and lifespan.

## I. INTRODUCTION

Aging exhibits remarkable diversity across species [1]. Short-lived animals such as flies or nematodes often display late-life mortality deceleration and plateaus [2, 3], whereas humans typically retain Gompertz-like acceleration until very late life [4–6]. At the other extreme, species such as certain turtles, rockfish, and the naked mole-rat exhibit *negligible senescence*, defined demographically as flat or nearly flat age-specific mortality rather than an absence of functional decline [7–9]. Yet across taxa, functional decline follows strikingly similar trajectories. Cardiorespiratory fitness, muscle mass, strength, and metabolic efficiency progressively diminish; recovery from perturbations slows; homeostatic responses weaken; and longitudinal variability in performance increases with age [10–12]. These shared features point to *universal principles* in aging dynamics, with distinct regimes—“normal” aging and negligible senescence—emerging from common underlying foundations [13, 14].

Despite these regularities, consensus on mechanisms remains elusive. More than 300 distinct theories of aging have been catalogued [15], with little agreement even among leading researchers. This fragmentation underscores the need for integrative frameworks that can bridge mechanistic detail with systems-level patterns [16]. In line with this view, the Dublin Longevity Declaration emphasized the urgency of developing quantitative syntheses to guide biomedical translation [17].

The existence of universal patterns suggests that aging may be captured by a small set of coarse-grained variables with clear biological meaning [18, 19]. In statistical physics, this property is known as *universality*: systems with very different microscopic constituents often display the same macroscopic behavior because their long-term dynamics are governed by only a few emergent variables. This reflects a key principle of complex systems—*separation of scales*—where the microscopic intricacy of physiology reduces, on aging-relevant timescales, to a handful of slow variables that dominate systems-level dynamics.

Building on these observations, we propose a minimal phenomenological model that makes the principle of universality explicit. In this framework, health trajectories and mortality are governed by three macroscopic variables: (1) a *dynamic response factor z*_0_, which captures stress responses and is characterized by a recovery time closely related to resilience; (2) an *entropic damage variable Z*, representing configurational entropy or information irreversibly lost during the aging process; and (3) a *regulatory noise strength D*_0_, quantifying the amplitude of stochastic physiological fluctuations. The central premise is that *z*_0_ acts as a fast variable whose stability margin is progressively eroded by the slow, linear accumulation of entropic damage *Z*, with *D*_0_ modulating the stochasticity of this coupled dynamics.

Within this framework, aging is defined as the coupled evolution of dynamic stress-response variables (*z*_0_) and the slow, largely irreversible accumulation of entropic damage (*Z*), modulated by intrinsic and extrinsic stochastic fluctuations (*D*_0_). This perspective constrains the achievable effect sizes of interventions. Therapies that act only on dynamic, age-dependent factors—corresponding to many of the established hallmarks of aging—are inherently limited in stable species such as humans, where resilience loss is driven primarily by damage accumulation. Larger and more durable gains would require reducing physiological noise or, more ambitiously, slowing the accumulation of damage itself, the latter offering the only route to shifting maximum lifespan. This classification of intervention levels thus provides a theory-grounded map from biological targets to realistic clinical outcomes.

In the sections that follow, we formalize this framework by linking resilience loss to thermodynamic irreversibility and critical dynamics. We then derive its empirical signatures—patterns expected in longitudinal aging data across species—and compare these predictions with observed trajectories. Finally, we discuss the implications for interpreting biomarkers, designing therapeutic strategies, and advancing the broader quest for effective interventions against aging.

## II. AGING AS AN EMERGENT PHENOMENON

Complex systems are hierarchically organized: at each scale, new variables and tools become relevant, reflecting behaviors that emerge from many interacting components. As P. W. Anderson famously observed, “More is different” [20]: the properties of large systems cannot be reduced to those of their parts. Classic illustrations include superconductivity (cooperative electron pairing), turbulence (nonlinear hydrodynamics), and phase transitions (macroscopic order with critical opalescence). Even fundamental thermodynamic quantities such as temperature are emergent—well defined only in the thermodynamic limit, while in small systems they fluctuate [21, 22].

A useful analogy is the second law of thermodynamics. At the microscopic level, physical laws are *time-reversal symmetric*, yet in macroscopic aggregates tiny uncertainties amplify through interactions, rendering reversal effectively impossible. This emergent asymmetry, expressed as entropy increase, produces the familiar “arrow of time” [23, 24]—and, as we argue below, an analogous arrow in aging biology.

The relationship between macro and micro is inherently asymmetric. Over long timescales, a few emergent variables suffice to capture system dynamics, while most microscopic degrees of freedom contribute only through statistical aggregates. In such regimes, collective variables dominate prediction and microstate details become inessential. This process of *coarse-graining* underlies *universality*: microscopically distinct systems can display the same macroscopic laws [25, 26]. Conversely, macroscopic constraints restrict the range of admissible microstates, exerting an inferential “top-down” influence, as formalized by Jaynes’s maximum-entropy principle [27, 28].

Because of this asymmetry, predicting macroscopic behavior directly from microscopic details is often intractable. Complex systems are more readily *observed* in certain states than *engineered* from the bottom up. This challenge is evident in drug discovery, where interventions act on molecular targets but the outcomes of interest—resilience, healthspan, and lifespan—emerge only at the organismal level. The mapping from micro to macro is inherently indirect and nonlinear, limiting the predictive power of purely reductionist approaches.

Aging provides an ideal domain in which to apply this framework, since the central challenge is to bridge microscopic mechanisms with macroscopic outcomes across widely separated timescales. It is a slow, organism-level process driven by myriad microscopic events. At the same time, species with radically different microscopic aging mechanisms exhibit convergent trajectories once coarse-grained. For example, extrachromosomal DNA circles drive aging in yeast but not in most other species [29], yet across taxa the phenomenology of aging follows strikingly similar patterns. Such convergence underscores the value of a coarse-grained, system-level description: it connects molecular mechanisms to long-term phenotypic change and provides a rational foundation for strategies to extend healthspan and longevity.

## III. PROBLEM STATEMENT AND EMPIRICAL TARGETS

Any theory of aging must account for a set of well-established empirical regularities. First, mortality trajectories follow near-universal patterns. In many species, mortality rates increase approximately exponentially with age over substantial adult ranges, consistent with the Gompertz law [4, 5]. The slope of this rise defines the mortality-rate doubling time. Deviations from strict Gompertz behavior are informative: some short-lived animals display late-life plateaus [2, 3], while species such as the naked mole-rat exhibit negligible senescence, showing little demographic acceleration despite clear molecular aging signatures [8, 9]. Humans are unusual in that Gompertz-like acceleration persists well beyond the average lifespan, with only weak signs of saturation at extreme ages [6]. In stark contrast, engineered systems typically fail with power-law hazards rather than exponential ones [30– That biology so consistently produces Gompertzian dynamics suggests a form of universality in aging that is absent from non-biological failure modes.

Second, aging is intrinsically stochastic. Even genetically identical individuals in controlled environments show broad variation in lifespan [10]. Across species, the variance of age-associated traits increases with age and scales inversely with lifespan [11, 33]. In humans, physiological indicators such as blood biomarkers display hyperbolic growth in variance, extrapolating toward a limit of ~ 120 years [33]. Rates of molecular drift also scale with longevity: somatic mutation and DNA methylation drift are lower in longer-lived species [34, 35], yet methylation drift persists even in negligible-senescence species such as the naked mole-rat [36].

A prominent molecular example comes from DNA methylation (DNAm) clocks. First-generation clocks predicted chronological age from CpG patterns with striking accuracy [37, 38], largely reflecting the stochastic accumulation of methylation variability—epigenetic drift [39–42]. Recent work formalized this insight into the concept of “stochastic clocks,” showing that dispersion alone accounts for 70–90% of predictive accuracy, while departures from strict stochasticity capture biologically informative processes [43–45].

Third, aging is constrained by cross-species scaling laws. Among mammals, lifespan and developmental time generally increase with body mass according to quarter-power scaling [46–48]. Conversely, rates of molecular and physiological change—including methylation drift, mutation accumulation, oxidative damage, and protein aggregation—tend to scale inversely with lifespan [49–51], reinforcing the tight coupling between damage accrual and longevity.

Fourth, aging is characterized by the low effective dimensionality of biomarker data. Analyses of genomic, molecular, and clinical datasets reveal that age-associated changes cluster, with only a few principal components capturing most of the variance [37, 38, 52, 53]. Measures of organismal performance and resilience—including frailty indices [54], maximal heart rate [55], lung function [56], and recovery times after sublethal stress [57]—decline approximately linearly with age at about one percent per year in humans, while interindividual variability increases.

A satisfactory theory of aging must therefore explain not only the average decline in function, but also the broadening of variance, the cross-species scaling laws, and the near-universal Gompertzian form of mortality together with its informative exceptions.

## IV. A MINIMAL AGING MODEL

To explain the empirical regularities of aging, a model must capture macroscopic trajectories without relying on intractable microscopic detail. Aging unfolds in living organisms—strongly interacting, open systems subject to continual noise. The underlying molecular composition and interaction networks are vast and only partly known, making a fully reductionist theory implausible. Yet interventions act precisely at this microscopic level, through small molecules, biologics, or cellular therapies. This creates a fundamental tension: we manipulate microlevel targets, but the outcomes of interest—resilience, healthspan, and lifespan—are emergent properties that exist only at the organismal scale.

Formally, the organismal state can be represented as a point **x** in a high-dimensional space, with each component *x*_*i*_ corresponding to a feature such as gene expression, CpG methylation, metabolite concentrations, or macromolecular integrity. Evolutionary constraints couple these features into coordinated processes, rendering many of the *x*_*i*_ highly correlated. As a result, the organismal state can be projected onto a lower-dimensional representation defined by collective “normal modes” *z*_*α*_, analogous to dynamical systems near equilibrium, in which subsets of features shift together when a mode is activated.

Each mode reflects a balance of three forces: (i) restoring tendencies that pull the system back toward homeostasis (resilience), (ii) persistent stresses such as smoking, malnutrition, or chronic inflammation, and (iii) stochastic fluctuations arising from both intrinsic sources (molecular errors, transcriptional bursts) and extrinsic ones (environmental variation). Mathematically, systems with these characteristics are described by Onsager’s linear nonequilibrium form, which captures how small perturbations relax toward equilibrium [58]:

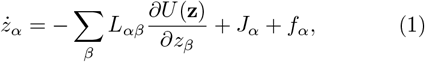

where *L*_*αβ*_ is the Onsager matrix, *J*_*α*_ represents persistent stresses, and *f*_*α*_ denotes stochastic noise. Near equilibrium, *∂U/∂z*_*α*_ ≈ Σ_*β*_ *H*_*αβ*_*z*_*β*_, with *H* the Hessian of the potential *U*. The stability matrix *K* = *LH* has eigenvalues *ε*_*α*_ that determine mode-specific recovery times: *ε*_*α*_ *>* 0 indicates stability, *ε*_*α*_ *<* 0 instability, and *ε*_*α*_ = 0 a bifurcation point.

While Eq. 1 describes such systems exactly, it is of limited practical use: the number of modes scales with the microscopic degrees of freedom, which in living systems is effectively infinite. Most of these modes cannot be measured, nor can their parameters be determined. Progress therefore requires *coarse-graining* based on three structural properties of aging systems.

First, modes with small eigenvalues evolve *slowly* and dominate long-timescale dynamics, whereas *fast* modes decay rapidly and contribute only smooth back-reaction forces. This separation of scales also explains the success of dimensionality-reduction methods such as PCA, which routinely extract slow, biologically meaningful modes [18, 59–61]. Thus, a small number of slow modes capture most of the organismal variance, while fast modes can be treated as *enslaved* —fluctuating around macroscopic averages set by the slow variables [62].

Second, the underlying free-energy landscape is rugged, with many local minima. Microscopic configuration changes—such as mutations, chromatin remodeling, methylation shifts, or protein modifications—correspond to rare transitions between these minima [52]. While individually infrequent, such events accumulate across billions of cells at appreciable rates. Because most departures from the evolutionarily optimized initial state reduce function, their aggregate effect is pathological. By the central limit theorem, only the total number of such transitions matters at macroscopic scales. We denote this cumulative burden as *Z*. In the simplest approximation, the individual events are independent, so *Z* follows a Poisson process and grows linearly in time. We refer to the growth rate *γ* as the damage accumulation rate (DAR).

Third, the stability of the system hinges on its slowest mode, the one with an eigenvalue *ε*_0_ closest to zero. Accumulated damage *Z* modifies this effective eigenvalue, and when *ε*_0_(*Z*) changes sign the system crosses from stability to instability. This motivates focusing on the critical mode *z*_0_, while other slow modes can be included as needed to describe subsystem failures underlying chronic diseases.

The formal derivation of a coarse-graining procedure based on these three structural properties is provided in Appendix A. Focusing on the critical mode and assuming weak couplings to other modes, the coarse-grained dynamics reduce to

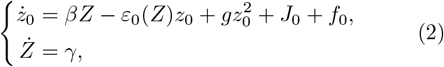

with *ε*_0_(*Z*) = *ε*_0_ − *β*^*′*^*Z*. Here *J*_0_ represents persistent stresses, *f*_0_ is a zero-mean white-noise term with autocorrelation ⟨*f*_0_(*t* + *τ*)*f*_0_(*t*)⟩ = 2*D*_0_*δ*(*τ*), and *γ* is the damage accumulation rate. The assumption of white noise is justified because *z*_0_ evolves on timescales much longer than the autocorrelation times of the fast modes that generate these fluctuations.

From this formulation, two observations immediately follow. First, the aggregation of microscopic transitions into *Z* illustrates emergence: organisms with different microscopic damage patterns but the same overall *Z* value are macroscopically indistinguishable on aging timescales.

Second, while individual modes exert negligible influence on *Z*, the reverse is not true: *Z* strongly modulates *z*_0_ and can drive *ε*_0_(*Z*) across zero, producing a bifurcation from stability to instability. In this sense, *Z* acts as a genuine macroscopic state variable of aging, one that emerges only in the thermodynamic limit.

## V. AGING PHENOMENOLOGY

Eq. 2, though minimal and dependent only on a few structural assumptions, captures a wide range of empirical phenomena. It reproduces critical slowing down, divergence of biomarker variance, Gompertzian mortality, and even explains systematic deviations from the Gompertz law. Crucially, its effective dynamics admit two qualitatively distinct behaviors set by system parameters. Nonlinear coupling to accumulated damage *Z* introduces a bifurcation: the system may sustain both a stable and an unstable fixed point, or collapse to a single unstable point (Fig. 1c). Biologically, these alternatives correspond to two fundamental regimes of aging that explain the diversity of trajectories across species and provide a natural basis for classification.

**FIG. 1.**
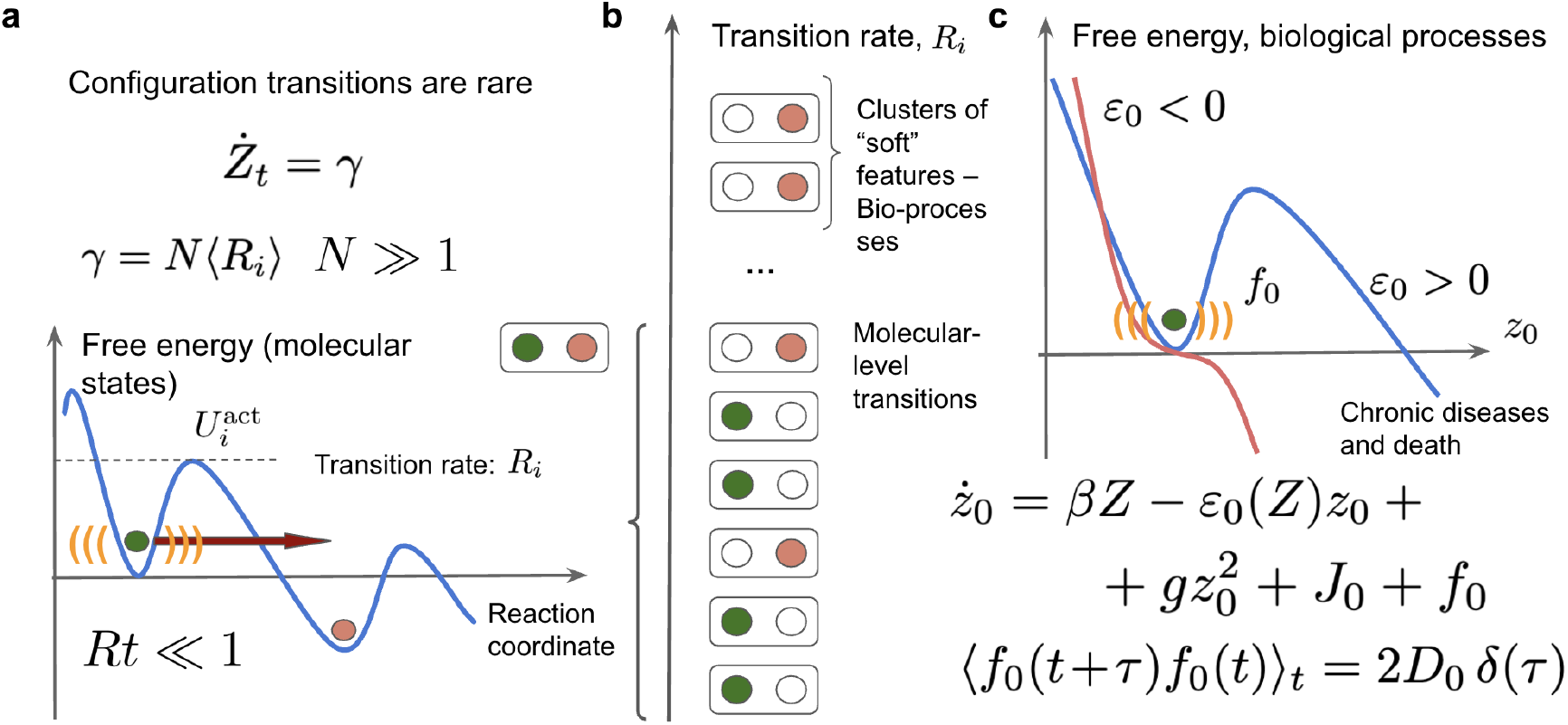
Schematic of the minimal aging model integrating entropic damage accumulation, dynamic stability loss, and mortality. **(a)** Rare configurational transitions between molecular states are thermally activated processes with high energy barriers. Although individual transitions occur infrequently (*R*_*i*_ ≪ 1), the cumulative damage rate *Ż*_*t*_ = *γ* = *N* ⟨*R*_*i*_⟩ can be substantial because of the large number of degrees of freedom (*N* ≫ 1). These events contribute to the slow, approximately linear accumulation of entropic damage *Z*, interpreted as generalized damage, configurational entropy, or irreversible information loss. **(b)** Transition rates arise from molecular events and clusters of coupled physiological features (“soft modes”) across hierarchical biological scales. Their aggregate behavior drives coarse-grained entropic dynamics at the organismal level. **(c)** Entropic damage *Z* modulates the stability of a critical dynamic variable *z*_0_, which governs resilience, stress response, and ultimately mortality risk. The effective free-energy potential for *z*_0_ depends on its recovery rate *ε*_0_(*Z*), which decreases with age. In the stable regime (*ε*_0_ *>* 0), fluctuations remain bounded; in the unstable regime (*ε*_0_ *<* 0), fluctuations diverge, leading to systemic collapse. The stochastic dynamics of *z*_0_ are governed by Langevin-type equations with nonlinear terms, persistent stress *J*_0_, and regulatory noise *f*_0_ characterized by ⟨*f*_0_(*t* + *τ*)*f*_0_(*t*)⟩ = 2*D*_0_*δ*(*τ*).

In the *stable regime*, a restoring fixed point keeps physiological fluctuations bounded, permitting recovery from perturbations, although resilience declines as *Z* grows. In the *unstable regime*, only an unstable point exists; trajectories diverge without restoring forces, driving rapid biomarker escalation and mortality acceleration. These regimes thus define distinct dynamical phases of aging.

In practice, the boundary between regimes is not fixed but shifts as the effective eigenvalue *ε*_0_(*Z*) declines with accumulated damage. Organisms may therefore begin life in the stable regime (*ε*_0_(*Z*) *>* 0) and cross into instability only once *Z* exceeds a critical threshold. The reverse trajectory is less evolutionarily plausible, since resilience during development and reproduction is strongly selected. Species can thus be classified by whether most of their lifespan unfolds under stability or instability: **Class I (unstable animals)**, in which instability is intrinsic from the outset, and **Class II (stable animals)**, in which resilience is initially maintained but ultimately lost through accumulated damage.

### A. Class I: “unstable” animals

In the unstable regime, the critical mode *z*_0_ begins life with a negative intrinsic recovery rate, *ε*_0_(*Z* = 0) *<* 0. This represents the extreme case where instability is present from the outset: physiological fluctuations are unconstrained, and resilience is absent from birth. In such systems, the slow accumulation of generalized damage *Z* plays only a secondary role, especially when the nonlinear couplings in Eqs. 2 are small. Mortality trajectories are therefore governed primarily by the intrinsic instability of *z*_0_. This regime, analyzed in detail by [18], follows a characteristic sequence of phases.

Early in life, stochastic fluctuations dominate, and the variance of *z*_0_ grows until it reaches 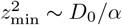, where *α* = |*ε*_0_|. Beyond this point, deterministic forces prevail, the effects of fluctuations may be neglected, and the mode variable grows exponentially along with its variance:

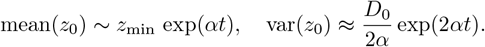

Eventually, the system crosses a nonlinear threshold at *z*_max_ = *α/g*, beyond which *z*_0_ diverges hyperbolically—a state incompatible with survival.

The average lifespan is therefore set by the thresholdcrossing time

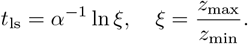

Because lifespan in unstable species is typically several times longer than the mortality-rate doubling time (MRDT ~ *α*^*−*1^), this implies *t*_ls_ ≫ *α*^*−*1^, hence *ξ* ≫ 1. Since *ξ* ~ 1*/g*, the nonlinearity *g* must be effectively small, explaining why most of life is spent in the exponential phase.

This prediction is supported by DNA methylation (DNAm) data in mice, which reveal two major signatures: (i) a linear component linked to damage accumulation (*Z*) and (ii) an exponential trend reflecting intrinsic instability along *z*_0_ [63]. A similar exponential component was recovered from longitudinal blood cell counts in the Mouse Phenome Database [64] using model-based fits of Eqs. (2) in [65], with doubling times matching empirical MRDTs - presicely as predicted by the model.

As derived previously in [18], survival probability takes the form *S*(*t*) = Erf(*ξe*^*−αt*^). Around the mean lifespan *t*_ls_, mortality follows an approximate Gompertz law:

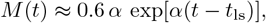

so the observed mortality doubling rate near midlife equals *α* = |*ε*_0_|. This implies that hazard curves around the average lifespan can be used to estimate *α* directly.

Because mortality depends strongly on *α* but only logarithmically on noise and nonlinearity through *ξ*, survival curves across conditions collapse under age rescaling by mean lifespan. This property has been validated experimentally in *C. elegans* [66].

The model also predicts systematic deviations from a pure Gompertz trajectory: shortly beyond the average lifespan, mortality acceleration slows and ultimately plateaus at a level close to the inverse MRDT, *M* (*t* → ∞) = *α*. Such deceleration and late-life plateaus have been observed in ultra-large cohorts and across multiple species [3, 18], including *C. elegans*, where the plateau matches the Gompertz slope measured earlier in life across strains with tenfold lifespan differences [67], and in mice from ITP studies, as analyzed in [65].

### B. Class II: “stable” animals

Unlike unstable species, “stable” animals begin life with positive intrinsic recovery rates, *ε*_0_(0) *>* 0. Their physiology can rebound from perturbations, maintaining a resilient equilibrium state. Early in life, fluctuations of the critical mode *z*_0_ are suppressed by strong restoring forces, and aging manifests primarily through the slow, linear accumulation of damage *Z*, which steadily erodes stability.

When *Z* = 0 and external stress is absent (*J*_0_ = 0), the free-energy landscape contains two fixed points: a stable minimum at *z*_0_ = 0 and an unstable maximum at *z*_0_ = *ε*_0_*/g*, which defines the protective barrier 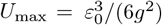 (Fig. 2). As damage accumulates, *ε*_0_(*Z*) declines approximately linearly, shrinking the distance between the fixed points and lowering the barrier. At the critical damage level *Z*_max_ = *γt*_max_, the recovery rate vanishes, *ε*_0_(*Z*_max_) = 0, and the barrier collapses in a saddle–node bifurcation. This transition defines the species-specific maximum lifespan,

**FIG. 2.**
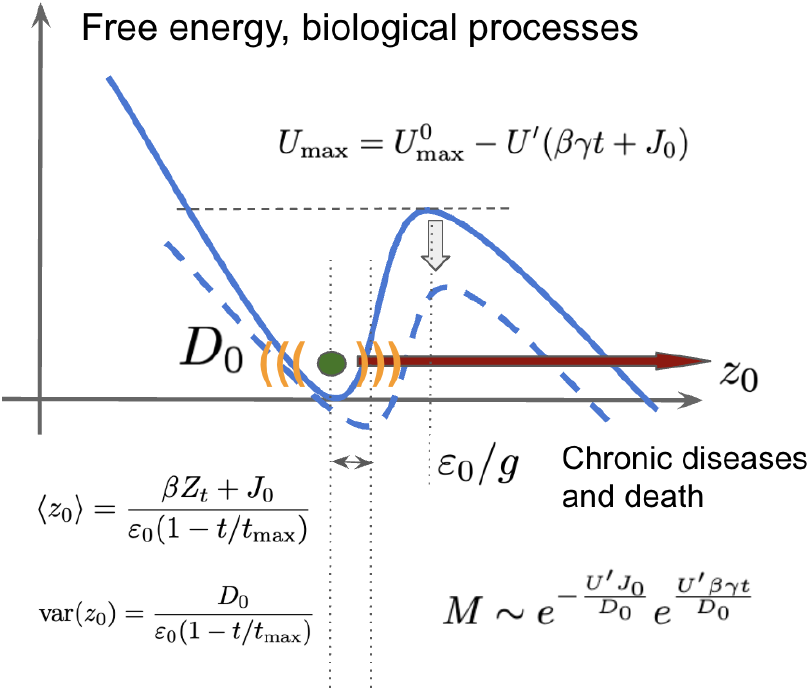
Stress and damage cumulatively erode the basin of attraction for the critical mode fluctuations. This erosion drives divergence in both the mean and variance of the critical mode while exponentially increasing the probability of escape transitions leading to death. The mechanism provides a natural explanation for the Gompertz mortality law, with the Gompertz exponent (inverse mortality rate doubling time, MRDT) proportional to the damage accumulation rate and inversely proportional to the magnitude of regulatory noise.

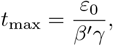

set by the initial stability margin, the coupling to damage, and the accumulation rate.

As *t* → *t*_max_, both the mean and variance of *z*_0_ diverge hyperbolically,

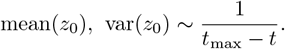

Persistent perturbations (*J*_0_) shift the mean but do not alter the variance, which is driven exclusively by *Z* ∝ *t*.

These predictions align closely with empirical data. Principal component analyses of mammalian DNA methylation consistently reveal (i) a linear component reflecting steady damage accumulation and (ii) a hyperbolically diverging component projecting to the species’ maximum lifespan (about 120 years in humans [52, 68]). Similar divergence has been documented in mortality-linked biomarkers including blood cell counts, physical activity, and composite metrics such as PhenoAge [12].

Death, however, typically occurs earlier, when stochastic fluctuations push *z*_0_ beyond the unstable point. Drawing on a foundational result from statistical mechanics, Kramers’ theory [69], mortality risk can be expressed as the probability of crossing an energy barrier that separates stable from unstable states:

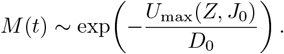

In the context of aging, this corresponds to the likelihood that resilience is lost and the organism transitions irreversibly from a functional to a failing state. Before *t*_max_, the barrier height declines approximately linearly with *Z, U*_max_ ≈ *U*_0_ − *U* ^*′*^*Z*, leading to exponential growth of mortality with age. This mechanism explains the ubiquity of the Gompertz law despite the fact that most types of molecular damage only accumulate linearly with age [70–72].

Note, that the Gompertz slope,

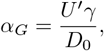

is directly proportional to the damage accrual rate and inversely proportional to noise strength.

A key implication is that the gap between average and maximum lifespan is determined by *D*_0_. In the noiseless limit (*D*_0_ → 0), barrier crossing would not occur until *t*_max_, eliminating the gap between average and maximum lifespan. In reality, stochastic fluctuations ensure that death typically occurs before *t*_max_, is reached, producing the observed distribution of lifespans. Empirical estimates of *D*_0_ from DNA methylation variance across mammals [68] confirm these model predictions. Thus, in stable animals, the combination of linear damage accumulation and stochastic barrier crossing explains both Gompertzian mortality and the separation of mean and maximum lifespan.

### C. Dynamic markers of aging and resilience

The model makes strong predictions for longitudinal data. The distinction between stable (*ε*_0_ *>* 0) and unstable (*ε*_0_ *<* 0) regimes can be revealed through the temporal autocorrelation function (TAC) of the critical mode *z*_0_:

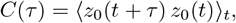

where ⟨·⟩_*t*_ denotes averaging first along individual trajectories and then across the cohort. Intuitively, the TAC measures how similar an organism’s present state remains to its past self over time. TAC analysis indicates not only whether a system is stable or unstable, but also quantifies the progressive decline of resilience as species approach the stability boundary.

In the stable regime, the TAC decays exponentially,

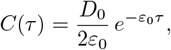

allowing *ε*_0_ to be estimated directly from longitudinal data. Its inverse, 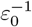, has the natural interpretation of a recovery time and thus provides a quantitative measure of resilience (see Appendix B for derivation).

Because *ε*_0_(*Z*) decreases with accumulated damage, longitudinal analyses of human cohorts should reveal a progressive slowing of autocorrelation rates with age—a hallmark of impending failure in the form of critical slowing down.

This pattern is indeed observed: autocorrelation times derived from complete blood counts and daily physical activity increase approximately linearly with age and extrapolate to zero around *t*_max_ ≈ 120 years [12]. This behavior is consistent with Eq. 2, in which variance diverges as resilience vanishes at the maximum lifespan. These data also suggest that *D*_0_ remains roughly ageindependent, since the observed increase in variance with age is fully explained by declining resilience without requiring an increase in stochastic fluctuations.

By contrast, in unstable species the TAC should show no age-related decay. Experimentally, In [65] we analyzed the Mouse Phenome Database [64] and found TACs essentially flat over 14–28 week intervals—far exceeding the ~ 4-week decay observed in humans. This absence of decay reflects the lack of a restoring force in the unstable regime and highlights the sharp difference in resilience dynamics across species.

Diverging patterns in TAC can therefore serve as dynamic markers of health and vulnerability. In [73], we showed that individuals with TAC values exceeding three months, indicative of lost dynamical stability, could be identified from longitudinal physical activity data. Importantly, their prevalence doubled approximately every seven years, consistent with Kramers’ barrier-crossing theory and the human mortality-rate doubling time of ~8 years.

The same framework also yields testable predictions for interventions. In unstable animals, perturbations of *z*_0_ produce persistent effects detectable in short-term studies, whereas in stable species TAC decays rapidly, so intervention effects fade once treatment ends. Smoking provides a familiar example: it elevates aging biomarkers, including epigenetic clocks, yet these changes reverse quickly after cessation [12, 73, 74]. More broadly, this difference in response to perturbations explains why shortlived models may overestimate the persistence of antiaging interventions in humans.

## VI. AGING AND THE SECOND LAW

Our phenomenological model distills the complexity of aging into the interplay of two macroscopic modes. The first is the dynamic variable *z*_0_, capturing leading organismal responses across physiological domains such as immune function, nutrient handling, and metabolic regulation. The second is the cumulative variable *Z*, representing damage that increases approximately linearly with age and progressively destabilizes critical physiological functions. In long-lived, “stable” species—whose average lifespan lies well below their maximum potential—this progressive accumulation of *Z* emerges as the dominant driver of aging.

Multiple datasets support the hypothesis that *Z* grows linearly with age. Principal component scores from large DNA methylation (DNAm) datasets—spanning mammalian species [75], humans [76], and mice [77]—show variances that increase proportionally with age, consistent with Poissonian stochastic processes expected if aging reflects the accumulation of many rare, independent molecular events [52, 68]. Single-cell DNAm analyses in aging mice [78] reinforce this view: CpGs contributing to the linear signature exhibit uniformly low mutual information, implying statistical independence of damage events [63]. In such a regime, both the mean and variance of *Z* increase linearly with age, while the number of accessible molecular configurations expands exponentially. Thus, *Z* can be naturally interpreted as configurational entropy and as a measure of cumulative information loss during aging.

In addition to being independent, site-specific transition (damage) rates in DNAm datasets are non-Gaussian and follow exponential-like distributions. Our recent analysis shows that this behavior is naturally explained if individual events correspond to thermally activated transitions over barriers whose heights follow a universal extreme-value (Gumbel) distribution [68]. Extreme-value theory predicts that dynamics are dominated by the highest barriers among a large set of possible transitions, implying that aging is driven by rare but consequential failures across otherwise redundant systems.

If *Z* reflects configurational entropy, its linear growth represents a direct manifestation of the second law of thermodynamics—an irreversible increase in entropy over time. This perspective clarifies why reversing accumulated damage is so difficult: it would require restoring an astronomical number of microscopic degrees of freedom to their initial, highly improbable configurations. In support of this irreversibility, heterochronic parabiosis in mice partially restored the dynamic (exponential) DNAm signature but left the entropic (linear) signature unchanged [63]. Thus, while the dynamic physiological response mode can be modulated or even reversed, the entropic mode appears fundamentally resistant to systemic rejuvenation.

The generalized damage tracked by *Z* is thus not just a biomarker but a fundamental state variable and the primary driver of aging in long-lived species (see also [79]). Its Poissonian kinetics, extreme-value rate distribution, and entropic nature place strong constraints on both theoretical models and realistic expectations for intervention outcomes. Moreover, because *Z* gradually modulates *z*_0_, it dictates population-level aging trajectories and sets the timing of resilience collapse.

## VII. INTERVENTION STRATEGIES: THREE LEVELS OF CONTROL

Our framework identifies three macroscopic variables as potential levers for intervention: the dynamic response factor *z*_0_, the cumulative entropic damage *Z*, and the regulatory noise strength *D*_0_. Each of these can, in principle, be modulated by biotechnological approaches or detected through phenotypic screening. In long-lived, stable species such as humans, aging is driven primarily by the gradual accumulation of *Z*, which progressively erodes resilience. Within this framework, intervention strategies naturally fall into three conceptual levels, differing in feasibility and expected impact (Table II).

**TABLE 1.** Comparison of aging characteristics in stable and unstable animals within the minimal model. Stable species exhibit bounded dynamics with resilience early in life and a finite maximum lifespan, while unstable species show divergence from birth and no intrinsic stability.

| Feature | Stable Animals | Unstable Animals |
| --- | --- | --- |
| Leading aging signatures | *Damage*: linear increase in mean and variance.<br>*Critical mode*: hyperbolic divergence of mean and variance approaching maximum lifespan. | *Damage*: linear increase in mean and variance.<br>*Critical mode*: exponential growth of mean and variance throughout life. |
| Maximum lifespan | Finite, determined by the initial stability margin and inversely proportional to the damage accumulation rate (DAR). | Absent in finite cohorts; survival limited only by stochastic fluctuations. |
| Resilience | Present early in life, progressively eroded; lost at the stability boundary near maximum lifespan. | Absent for the critical mode from birth; trajectories diverge without restoring forces. |
| Gompertz exponent $\alpha_G$ | Non-universal; proportional to DAR and inversely proportional to regulatory noise $D$. | Equals the intrinsic instability rate $\varepsilon_0$. |
| Mortality acceleration | Exponential increase up to maximum lifespan, after which mortality diverges. | Exponential increase until the mean lifespan, followed by a late-life plateau. |
| Temporal autocorrelation function (TAC) | Exponential decay, with decay rate decreasing linearly with age (slowing recovery). | No decay; TAC remains flat over life, reflecting absent resilience. |
| Effects of drugs targeting the critical mode ($z_0$) – hallmarks of aging | Transient: reduce biomarkers and risk only during treatment; effects dissipate after cessation. | Persistent: short interventions can produce lasting effects on risk and biomarkers. |
| Effects of drugs targeting noise ($D_0$) | Strong impact: lowering $D_0$ narrows the lifespan distribution and “squares” the survival curve. | Weak or negligible effects, as dynamics are dominated by instability. |

**TABLE 2.**
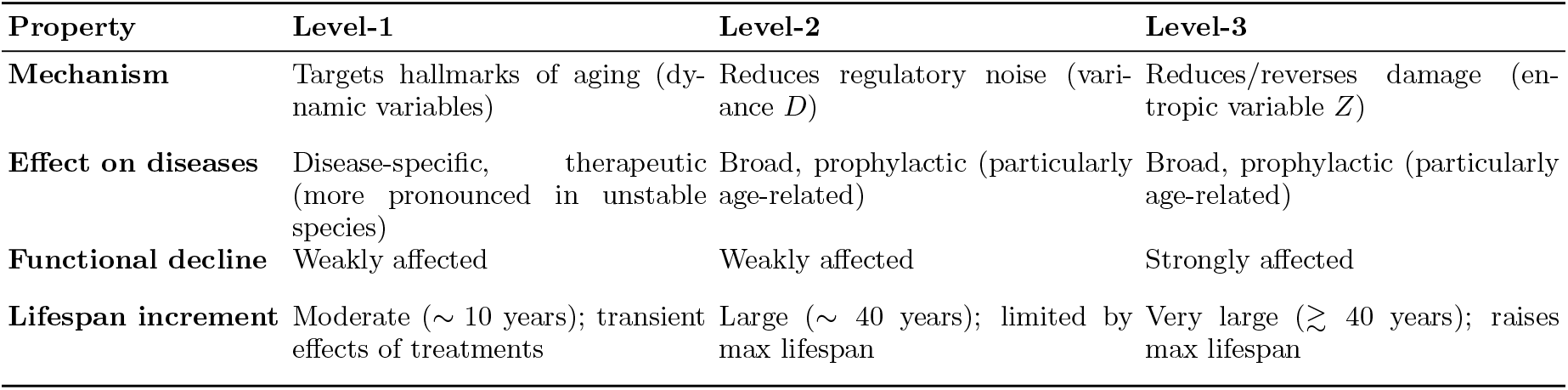
Comparison of three classes of anti-aging interventions in long lived species.

### Level 1: Modulating dynamic response factors (*z*_0_)

Level-1 strategies act on physiological response systems that become dysregulated with age. In our framework, these correspond to the dynamic response mode *z*_0_, though in practice multiple slow response modes may contribute. Dysregulation and resilience loss in these modes manifest as chronic diseases such as diabetes, cardiovascular disease, and cancer. Most current therapeutic approaches—including caloric restriction mimetics, senolytics, telomerase activators, and cellular rejuvenation strategies—fall into this category.

Interventions targeting dynamic response modes are especially effective in short-lived organisms, where the critical dynamics of the most unstable mode are tightly coupled to lifespan. In humans, however, their primary benefit lies in preventing or delaying specific chronic diseases with high population burden. For example, type 2 diabetes alone can reduce life expectancy by up to eight years depending on age at onset, underscoring the value of metabolic optimization within this category.

### Level 2: Reducing physiological noise (*D*_0_)

Level-2 strategies aim to reduce physiological noise (*D*_0_)—the random fluctuations that can push organisms beyond their limits of resilience. In our framework, lowering *D*_0_ decreases the probability of stochastic events that tip individuals into frailty or disease. Importantly, reducing physiological noise does not slow the accumulation of *Z* and therefore cannot extend the maximum lifespan, the age at which resilience is fully eroded by damage.

Level-2 interventions could, however, substantially reduce the risk of premature physiological collapse, effectively rectangularizing survival curves by compressing morbidity without altering the baseline rate of decline. Extrapolating from demographic data, such strategies could extend healthy lifespan by 30–40 years, bridging the gap between today’s average lifespan (70–80 years) and the maximum observed limit (~120–150 years).

### Level 3: Controlling entropic damage (*Z*)

Level-3 strategies address the cumulative entropic damage *Z*, the fundamental driver of aging in long-lived species. Whereas Level-1 approaches modulate dysregulated systems and Level-2 approaches reduce stochastic noise, Level-3 interventions seek to slow the accumulation of *Z* or mitigate its functional consequences. Because *Z* reflects configurational entropy, it is unlikely to be reversed by current technologies.

Nonetheless, several conceptual avenues exist: molecular repair technologies, clearance of irreversibly damaged cells or macromolecules, organelle replacement, genome and epigenome editing, or large-scale cell and organ replacement therapies. The most ambitious strategies would ultimately aim to arrest the linear growth of *Z* altogether—an achievement that would not only extend lifespan but also fundamentally reshape the trajectory of functional decline, shifting the maximum lifespan itself.

Realizing such interventions will require transformative enabling advances, including reliable biomarkers to track *Z*, scalable platforms for tissue and organ repair, and rigorous validation in long-lived animal models. While technically daunting, our framework shows that these strategies are scientifically coherent and provides a structured basis for defining the innovations needed to make them feasible.

## VIII. DISCUSSION

Our minimal theoretical framework provides a semiquantitative theory of aging that advances the field in two key ways. First, it demonstrates how diverse hallmarks and microscopic damage processes—collagen cross-linking, epigenetic drift, mitochondrial dysfunction, cellular senescence, and others—can be coarse-grained into a single entropic variable *Z*, naturally interpreted as configurational entropy. Processes that are effectively irreversible, such as somatic mutations or chromosomal abnormalities, contribute directly to the age-dependent accumulation of *Z*. By contrast, partially reversible processes—including proteostasis imbalance, accumulation of senescent cells, or chronic inflammation—primarily perturb *z*_0_, the dynamic response mode(s). This coarsegrained perspective aligns with and extends the concept of the *deleteriome* [80], which emphasizes the cumulative burden of deleterious changes as the central driver of aging.

Second, the framework explains how stochastic microscopic events, by progressively renormalizing slow physiological modes, generate universal patterns of resilience loss, morbidity, and mortality across species. This resolves a longstanding paradox in the field. Damage-accumulation theories have a long history, with more than 300 proposals catalogued [15]. Among the most influential was the Free Radical Theory of Aging (FRTA) [81], later refined into the Mitochondrial Free Radical Theory (MFRTA) [82], which emphasized mtDNA damage as both source and target of reactive oxygen species, creating a vicious cycle of exponential damage accumulation. While early studies supported a central role for oxidative stress [83–86], later work challenged this view: antioxidant trials often failed [87], some long-lived mutants exhibited elevated ROS [88], and mutator mice with high mtDNA mutation rates aged prematurely without dramatic ROS amplification [89]. Today, mitochondria remain central to aging biology, but oxidative stress is recognized as one contributor among many rather than a single universal cause.

In our synthesis, aging is not the consequence of a single causal lesion but emerges as a generic dynamical instability. Small stochastic perturbations are absorbed by stress responses and rare residual damaging events lead to build up of the entropic load *Z*, progressively destabilizing slow resilience modes until critical thresholds are crossed. Microscopic processes—oxidative lesions, collagen cross-links, epigenetic drift, chronic inflammation, mitochondrial damage—are individually rare, but in aggregate renormalize the dynamics of system-level variables. Their coarse-grained effect is largely mechanism-independent, yet naturally produces the exponential rise in mortality risk described by the Gompertz law.

Aging arises from the collective imprint of molecular events, whether through the critical instability of specific dynamic factors or the cumulative accumulation of entropic damage. In both cases, it emerges as a quantifiable causal process in its own right. This perspective accounts for the near-universal Gompertzian mortality pattern, explains why diverse hallmarks converge on common trajectories of functional decline, and provides a theoretical basis for interpreting epigenetic and other molecular aging clocks as statistical readouts of entropic load together with signatures of resilience loss.

In this way, the framework clarifies how to interpret epigenetic aging clocks. In our model, aging signals arise from two coupled sources: microscopic configurational transitions in fast modes and gradual erosion of stability in organism-level resilience (slow modes). First-generation clocks, such as the Horvath [37] and Hannum [38] models, are regression-based predictors that combine variability from both sources, but predominantly reflect the entropic component of epigenetic drift [42]. Stochastic clocks similarly capture this composite variability and are likewise dominated by the entropic contribution [43].

By contrast, mortality- and morbidity-trained clocks, including PhenoAge [90], GrimAge [91], and DunedinPACE [92], place greater weight on slow-mode components (*z*_0_) that are directly linked to functional decline and morbidity. Both classes therefore provide complementary windows into the same underlying dynamics, albeit with different weightings of entropic and dynamic contributions. Yet neither can fully disentangle these signals, since stochastic clocks also capture slow-mode responses to cumulative entropic load, while disease-trained clocks remain sensitive to resilience loss that is itself proportional to *Z*.

Our framework extends the classical Strehler–Mildvan (SM) theory [93] by mapping the SM concept of a “vitality deficit” onto the cumulative damage *Z*, identifying *z*_0_ as the reaction coordinate underlying Gompertzian mortality, and assigning *D*_0_ a temperature-like role – all identifiable from molecular level data. In this way, parameters that were purely empirical in the original SM theory acquire clear physical and biological meaning.

Related mechanistic models, including those of Alon and colleagues [19, 94, 95], emphasize subsystem bifurcations or distinct “ballistic” versus “quasi-steady” regimes. While such models highlight important biological mechanisms and, like ours, derive Gompertzian mortality via a Langevin process undergoing bifurcation, they differ in what drives the transition. Typically, these models invoke strong nonlinearities (e.g., saturation of repair pathways) to generate regime switching. By contrast, our framework requires only weak nonlinearities: as the recovery term *ε*_0_ approaches zero, even small couplings to *Z* qualitatively shift system behavior.

Our approach resonates with broader perspectives in complexity science, such as Wolfram’s “computational universe” [96], where simple microscopic rules generate robust macroscopic regularities. By invoking the ideas of emergence through coarse-graining and highlighting universalities, the model establishes a conceptual bridge to statistical physics. Physiological dynamics evolve near a critical point separating stable and unstable regimes, echoing the notion of systems operating at the “edge of chaos,” where order and adaptability are balanced [97]. Renormalization Group arguments [98] reinforce this perspective: in large systems, nonlinear couplings renormalize to weak effective values, consistent with both the simplicity of the effective equations and empirical evidence. In particular, unstable animals display extended exponential “ballistic” phases of biomarker change (see Section V), illustrating how weak nonlinearities combined with criticality may shape aging trajectories. Thus, our universality-based framework generalizes mechanistic theories by embedding them within a broader class of critical aging dynamics.

Negligible senescence represents an important edge case: species in which demographic aging—the Gompertzian acceleration of mortality—is absent or delayed far beyond the lifespan expected for animals of comparable body mass. The naked mole-rat provides the best-studied example: despite clear molecular aging signatures, including progressive epigenetic drift and entropy growth consistent with *Z* [99], demographic cohorts show no Gompertzian acceleration within the observed age range [100, 101].

In our framework, negligible senescence arises when the coupling between *Z* and *z*_0_ is weak, producing mortality rates that remain nearly constant with a vanishingly small Gompertz slope, or when *D*_0_ is exceptionally low, suppressing age-dependent mortality until very near the maximum lifespan. Thus, longitudinal studies of negligible-senescence species should reveal one of these two signatures, providing a critical empirical test of the framework.

Finally, by distinguishing the stochastic but reversible dynamics of *z*_0_ from the nearly deterministic, linear accumulation of *Z*, the framework embeds aging within the second law of thermodynamics. The statistical independence of damage events implies exponential growth of accessible configurational states, giving *Z* a natural interpretation as configurational entropy and cumulative information loss. This perspective highlights the fundamental challenges on the way to reverse of aging in long-lived species.

## IX. CONCLUSION

We have introduced a minimal theoretical framework in which aging dynamics are governed by three macroscopic variables: the dynamic response factor (*z*_0_), cumulative entropic damage (*Z*), and stochastic noise (*D*_0_). Together these variables capture how organisms balance stress responses, irreversible deterioration, and random fluctuations. This abstraction unifies diverse hallmarks of aging, explains universal demographic laws such as Gompertz scaling, and links directly to empirical biomarkers including DNA methylation clocks and physiological resilience measures.

By treating aging as the outcome of the interplay between physiological stress responses and cumulative damage—the aggregate effect of countless molecular-level configurational transitions—our framework shifts focus from isolated mechanisms to universal variables with predictive power. It also provides a principled taxonomy of interventions: modulation of stress-response dynamics (*z*_0_), reduction of physiological noise (*D*_0_), and control of entropic damage (*Z*).

In this view, aging is neither reducible to a single hallmark nor attributable to one microscopic mechanism. It is a collective, thermodynamically irreversible process that emerges from the aggregation of many stochastic events. Yet while *Z* embodies irreversibility, its rate of growth remains modifiable. Interventions that slow damage accrual could shift human aging trajectories in fundamental ways, potentially extending lifespan beyond current natural limits.

By reducing aging to a minimal set of measurable variables, our framework offers both a unifying language for theory and a roadmap for the rational design of interventions targeting the aging process.

## ACKNOWLEDGMENTS

We are deeply grateful to Pasquale Di Cesare for many stimulating discussions and thoughtful suggestions throughout the development of this work. His engagement with both the manuscript and the underlying ideas greatly improved their clarity and scope. We thank Vadim Gladyshev for his constructive feedback on earlier versions of this manuscript, including thoughtful comments on negligible senescence, conditional reversibility of damage, and the role of hallmarks, which helped sharpen the conceptual framing. We also thank Uri Alon for insightful discussions and for sharing his complementary theoretical perspective on universal patterns of aging, disease incidence, and functional decline, which enriched the comparative analysis presented here. Their input reflects the spirit of collaborative inquiry that is essential for advancing a unified understanding of aging.

## Notes

### Competing Interest Statement

The authors have declared no competing interest.

### Summary of Updates

a misprint in the abstract - no findings changed or removed

